# Effect of copper mill waste material on benthic invertebrates and zooplankton diversity and abundance in Lake Superior beaches

**DOI:** 10.1101/2024.03.20.585947

**Authors:** James H Larson, Michael R Lowe, Sean W Bailey, Amanda H Bell, Danielle M Cleveland

## Abstract

From 1900 to 1932 a copper (Cu) mill operated near Gay, Michigan, along the eastern shore of the Keweenaw Peninsula (Lake Superior, Michigan) and discharged waste material (stamp sands [SS]) to a nearby beach. These SS escaped containment structures and have been redeposited by wave action along the beaches in northern Grand Traverse Bay and onto Buffalo Reef, an important spawning area for native fish. Newly hatched fish move into nearby beach habitats where they grow during their first summer. Juvenile fish initially consume zooplankton before switching to benthic invertebrates once they are large enough. SS contain metals (especially Cu) that are toxic to many invertebrate taxa, and studies have observed few benthic taxa in areas covered by SS. We sampled the invertebrate community from four Lake Whitefish nursery areas: one near Buffalo Reef with high SS, one south of the Traverse River with moderate SS, one in nearby Little Traverse Bay with little SS, and a beach ∼58 km away with no SS (Big Bay). We also resampled the benthos at sites that had been sampled as part of an earlier Grand Traverse Bay study. Buffalo Reef (high SS) had fewer benthic taxa, and less density of several taxa than Little Traverse Bay (little SS), especially benthic copepods. All beaches had comparable zooplankton diversity, but the abundance was ∼2 orders of magnitude lower at Buffalo Reef (high SS) than other beaches. Cu and several other metals were elevated at beaches with more SS. We found support for associations between benthic density and diversity with depth (positive effect) and Cu concentration (negative effect). Cu concentration was a better predictor of declines in benthic invertebrate abundance and diversity than SS. We also observed that the relationship between Cu concentration and SS was non-linear, and highly variable. For example, 149 mg Cu/kg dry weight sediment is a consensus toxicity threshold used in the literature, but the prediction interval around that concentration from our model is 26-851 mg Cu/kg dry weight. A better predictive model of this relationship would be beneficial to develop to understand what level of SS reduction would prevent Cu impacts on invertebrates.

## Introduction

Historical copper mining in the watersheds surrounding Lake Superior was done by bringing ore-bearing rock to stamp mills, that crushed the rock into a consistent size for copper removal. These ‘stamp sands’ (SS) were moved to nearshore structures that have since failed and contaminated many nearshore ecosystems (Kerfoot and Nriagu 1999, Jeong et al. 1999). One location where SS were deposited was on the eastern shoreline of the Keweenaw Peninsula, Lake Superior near Gay, Michigan. Between 1900 and 1932, approximately 22.8 million metric tons (Mt) of SS was discharged to the beach near the mill (Kerfoot et al. 2012). Of the original 22.8 Mt, an estimated 11 Mt of SS remains on the shoreline, and the remainder (minus approximately 1.4 Mt removed by the local road commissions for use as traction sand) has spilled out into Grand Traverse Bay and into Buffalo Reef (Kerfoot et al. 2019, 2021).

Buffalo Reef is a 2,200-acre cobble and bedrock formation that is situated 5 km south of the original pile of SS in 3–30 m of water and less than 100 m from the stamp sand-covered beach. The reef has long been recognized as an important spawning habitat for Lake Trout (*Salvelinus namaycush*) and Lake Whitefish (*Coregonus clupeaformis*) (Goodyear et al. 1982, Chiriboga and Mattes 2008). Buffalo Reef supports valuable tribal, commercial, and recreational fisheries, and the waters within 80 km of Buffalo Reef historically produced more than half of the Lake Trout and Lake Whitefish harvested in western Lake Superior’s 1842 Treaty waters (Mattes 2020). However, that level of harvest began to decline significantly in the early 2000s, coincident with when Tribal fishers first expressed concern about SS drifting onto the reef and affecting critical habitats (Chiriboga and Mattes 2008). Early work on Buffalo Reef determined that not only do stamp sands contain copper (Cu) in concentrations that are toxic to cold-water fishes (MDEQ 2012), but also that >25% of the reef’s surface was already covered with SS in 2009 (Kerfoot et al. 2019). By 2015, the extent of the reef covered by SS had increased to 35% (Kerfoot et al. 2019), and hydrodynamic models predicted that 60% of the reef would be inundated by SS by 2026 if no management action was taken (Hayter et al. 2015). More recently, biologists at the Great Lakes Indian Fish & Wildlife Commission (GLIFWC) have noted an increasing presence of SS in the natural sand beach along the lower half of Grand Traverse Bay (Mattes, written comm., 2020). This suggests that the drifting SS have circumvented the breakwall that protects the entrance to the Traverse River harbor.

As a result, Buffalo Reef and the Gay SS pile have become the focus of a large-scale mitigation and protection effort lead by the Buffalo Reef Task Force (BRTF). The BRTF, which is composed of representatives from multiple Tribal, state, and federal regulatory and science agencies, has developed a long-term plan for removing the SS from the shoreline (BRTF 2023). In turn, the BRTF has directed a core ecology team to evaluate the potential ecological outcomes for each of the task force’s proposed management alternatives, including ‘no action.’ However, few biological investigations have been conducted in relation to Buffalo Reef, making it difficult to assess the impacts of drifting SS on important fish habitats and fish production. Much of the available information is derived from either toxicity testing by the State of Michigan or from routine monitoring and assessment efforts by GLIFWC. Those data show (1) reduced survival and growth of Amphipoda and Cladocera exposed to SS elutriates (MDEQ 2006), (2) declining numbers of spawning Lake Trout and Lake Whitefish in recent years (Mattes 2020), (3) absence of young-of-year (YOY) Lake Whitefish from the SS-covered beaches adjacent to Buffalo Reef (Premo 2008), and (4) sharp declines in YOY Lake Whitefish abundance in the juvenile nursery habitat (i.e., natural sand beach in the southern portion of Grand Traverse Bay) since 2008.

The mechanisms driving these observed changes are poorly understood because SS represent an unusual environmental contaminant due to a capacity to exert both chemical and physical changes to the ecosystem. The drifting SS have physically altered the surface of the reef and smothered its nearshore area (Kerfoot et al. 2021) where Lake Whitefish once spawned (Chiriboga and Mattes 2008). Even in areas where the surface of the reef is not inundated, SS have likely settled into the interstitial spaces that are critical to the survival of the early life-stages of Lake Whitefish (Ebener et al. 2021). Moreover, *in situ* conditions within the reef are difficult to assess, due to the water depths and temporal variations in SS presence (e.g., drifting, leaching, weathering).

Natural mortality is generally high for Lake Whitefish embryos in relatively pristine habitats (27–100%; Taylor et al. 1987, Freeberg et al. 1990), and is often linked to winter temperatures, ice formation, and availability of larval prey items. Many common prey items for larval and juvenile fishes (Claramunt et al. 2010, Pothoven et al. 2014) can be sensitive to metals, resulting in effects such as reduced populations or extirpation. Zooplankton and benthic invertebrates can also take up metals via ingestion of contaminated algae, sediment, periphyton, and other detritus, which creates another exposure pathway for fish. Thus, the movement of SS into the shallow nursery area south of the Traverse River could also harm fish communities by threatening their prey resources (Kerfoot et al. 2012, 2019, 2021). This is especially concerning for Lake Whitefish because juveniles use these habitats for the first five months of life (Claramunt et al. 2010).

In a pair of studies, Kerfoot et al. (2019, 2021) linked the presence of SS to a biological response at Buffalo Reef, summarizing several years of benthic sampling data to demonstrate that both taxonomic diversity and abundance of benthic macroinvertebrates are negatively affected by SS. Kerfoot et al. (2019, 2021) found that most taxa were absent in the areas where SS have almost completely covered native sands and identified a strong, direct relationship between SS percentage and both benthic density and diversity. Here, we sought to add to this work by quantifying the impact of stamp sands on benthic and zooplankton communities at beaches that varied in their proximity to the Gay stamp sand pile. We also resampled a subset of sites sampled by Kerfoot et al. (2021) to assess recent changes in both SS concentration and benthic community composition. These data were used to model the relationship between the benthic community and the proportion of stamp sands in the substrate.

## Methods

### Study Area

Our study area consisted of four beaches spread across Keweenaw Bay, Lake Superior. Each beach had varying stamp sand presence, based on previous mapping of the stamp sands distribution by Kerfoot et al. 2021 (Figure 1). Beaches were the focus of this study because these are important foraging areas for juvenile fish spawned on the nearby reefs. Two beaches were located in Grand Traverse Bay (Lake Superior), one adjacent to Buffalo Reef with high SS (BUR-H) and one south of the harbor breakwall at the Traverse River with moderate SS (Grand Traverse Bay beach; GTB-M) (Figure 1A). The breakwall was installed in 1950 to provide a safe harbor for boats; however, it also reduced the southeastern movement of the SS and, until recently, prevented SS intrusion at GTB-M. The drifting SS eventually overtopped the wall and now low amounts (<15% in the nearshore environment) of SS are found in some areas of GTB-M (Kerfoot et al. 2021). Conceptually, BUR-H represents a site that has been substantially inundated by SS, whereas GTB-M has experienced comparatively fewer SS to date. In addition to the beach sampling areas in Grand Traverse Bay, we resampled 16 sites (Figure 1) that had previously been reported (RES) in Kerfoot et al. (2019).

**Figure 1.**
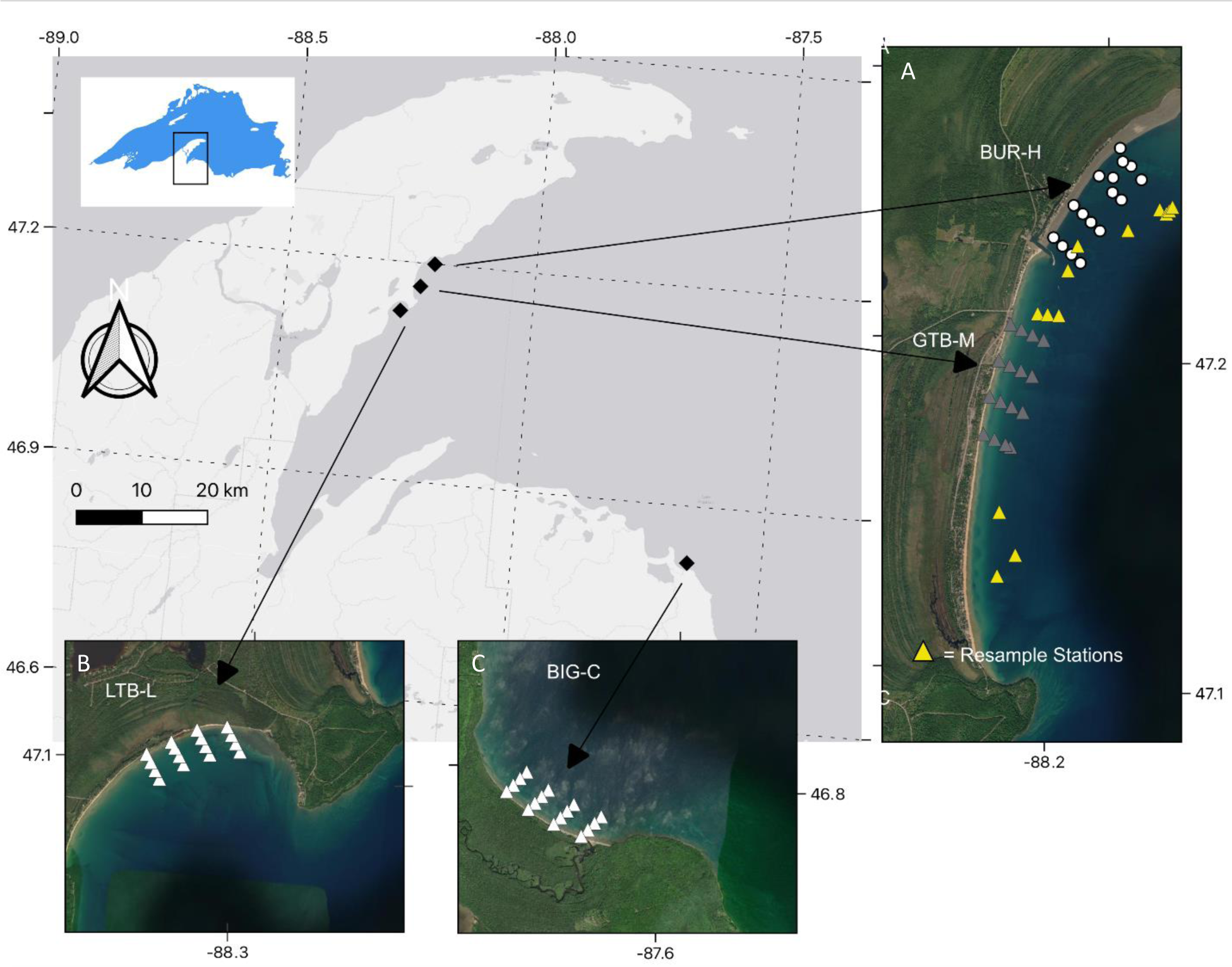
Map showing the locations of the study sites off Michigan’s Keweenaw Peninsula in Lake Superior, Michigan in 2021. Sampling transects and locations of Petite Ponar grabs at Buffalo Reef (BUR-H; white circles), Grand Traverse Bay (GTB-M; grey triangles), Little Traverse Bay (LTB-L; white triangles), (BIG-C; white triangles), and sites previously reported by Kerfoot et al. (2019; Resample stations; yellow triangles). Base map from Esri and its licensors, copyright 2023.

Two other sites were sampled with low or no SS influence. Continuing southeast along the Keweenaw Peninsula is a beach in Little Traverse Bay that experiences very low SS influence (LTB-L). We are unaware of previously published mapping of SS at this location, but given the distance of LTB-L from Gay, Michigan, and the orientation of the shoreline (i.e., LTB-L is shielded from SS migration by Traverse Point to the north/east; Figure 1B), this site is likely to have low SS relative to BUR-H. A control site was the beach at Big Bay (BIG-C), which is approximately 58 km from the original SS pile, on the opposite side of the Keweenaw Bay near Big Bay, Michigan. The BIG-C site is unlikely to receive SS from the pile at Gay, Michigan, due to the overall distance separating the two locations. A nickel-copper mine is currently operating in the BIG-C watershed, but it has a relatively small footprint (surface operations: 53 ha) and does not produce SS; ore is hauled offsite and farther inland for processing. There are no other records of SS on the Lake Superior shoreline along the southern edge of Keweenaw Bay or in the vicinity of BIG-C. However, like other Lake Superior beaches, other industries (e.g., logging and production of wood products) have historically been very active in the BIG area and may have served as a source for other, untested contaminants. This site also has a different orientation than the other sites, ‘facing’ north/northwest (N/NW) (out of Keweenaw Bay), whereas the other beaches face south/southwest (S/SW) (the opposite side of Keweenaw Bay). As a result, the beach at BIG-C is more exposed to wind and wave action from the main fetch of Lake Superior.

### Sediment and macroinvertebrate sample collection

At each beach, we collected sediments and benthic macroinvertebrates at 16 stations arrayed along four transects that were perpendicular to the shoreline, across a range of water depths (Figure 1B, C, and D). Sediment and invertebrates were collected between August 17 and August 19, 2021, for BUR-H, GTB-M, LTB-L and RES samples, and on August 23, 2021, for BIG-C. Samples were collected with a Petite Ponar grab sampler (Wildco; Powers and Robertson 1967) and separated into sediment and macroinvertebrate fractions in the field. The Ponar sampler was thoroughly cleaned between each sampling site. Each sediment sample was mixed well on the boat, and a subsample of sediment (approximately 65 g) was collected and split for grain size analysis, including estimation of percent SS, and analyses of total recoverable metals. Subsamples were frozen until further processing. The remaining portions of the grab sediments were then sieved (500-µm mesh) to remove small debris and capture benthic invertebrates. Material not passing through the sieve was decanted into plastic bottles filled with 10% neutral buffered formalin (NBF) and a Rose Bengal dye for macroinvertebrate identification and quantification. Water temperature (°C), dissolved oxygen (mg/L), and pH were recorded at each Ponar grab site (just above the water-sediment interface) prior to sample collection.

### Sediment processing

Sediment sub-samples for SS determinations were taken to the U.S. Geological Survey’s (USGS) Hammond Bay Biological Station, thawed in the laboratory, oven-dried to a constant temperature at 65 °C, and then passed through a series of sieves (4-mm, 2-mm, 500-µm, 250-µm, 125-µm, and 63-µm) for grain size analysis. To estimate the amount of SS in each sediment, three replicate samples were collected by pressing double-sided tape into the surface of the dried sample (after thorough mixing). This tape was then affixed to a glass microscope slide with an etched, 9 x 9, labelled grid (each grid cell was 1.0 mm x 1.0 mm); slides were imaged at 20x magnification using a 12-megapixel CMOS camera (AmScope) fixed to a compound microscope (Zeiss Primo Star). Four cells from each grid were randomly selected and the number of stramp sand and natural quartz sand grains were enumerated in each of the cells by two different individuals.

Sub-samples of frozen sediments for analyses of total recoverable metals were sent to the USGS Columbia Environmental Research Center in Columbia, Missouri. Arsenic (As), cadmium (Cd), chromium (Cr), cobalt (Co), copper (Cu), lead (Pb), iron (Fe), manganese (Mn), nickel (Ni), selenium (Se), thallium (Tl), and zinc (Zn) in the sediments were quantified using inductively coupled plasma-mass spectrometry (PerkinElmer Nexion 2000; similar to U.S. Environmental Protection Agency [US EPA] 6020B; US EPA 2014) following microwave-assisted digestion in high purity nitric acid and 30% hydrogen peroxide (similar to US EPA method 3050B; US EPA 1996). This digestion method accesses elemental concentrations that could become biologically or environmentally available; it is not a total digestion method. All sediment samples were lyophilized prior to digestion, and elemental concentrations were determined in units of milligram per kilogram on a dry weight basis (mg/kg dw). Quality control (QC) measures included the use of at least four NIST-traceable calibration standards, second-source initial and continuing calibration standards and blanks, and laboratory control standards, as well as method digestion blanks, digestion triplicates and spikes, and the use of certified reference materials having a soil or sediment matrix; all QC results were within acceptable limits for the method. The limits of quantification (LOQs) were the following (mg/kg dw): As, 0.04– 0.1; Cd, 0.02–0.04; Co, 0.02–0.03; Cr, 0.04; Cu, 0.1–0.2; Fe, 1–2; Mn, 0.2–0.4; Ni, 0.1–0.2; Pb, 0.02; Se, 0.1; Tl, 0.02–0.03; and Zn, 0.3–0.4. Concentrations of Cr, Mn, Fe, Co, Ni, Cu, Zn, As, and Pb in all 79 sediments were above their respective LOQs. There were 31 sediments having Se concentrations <LOQ (39%) and 38 Cd concentrations <LOQ (48%); all 79 sediments had Tl concentrations <LOQ (100%).

### Macroinvertebrate processing

Benthic macroinvertebrate samples were processed at the USGS Upper Midwest Environmental Sciences Center in La Crosse, Wisconsin. The NBF/Rose Bengal dye solution was decanted and replaced with 95% ethanol to facilitate processing and identification. Using a zoom stereo microscope (7–70x magnification; Olympus SZH10), invertebrates were sorted and identified to at least the level of taxonomic classification used in Kerfoot et al. (2019). In most cases, this was to the Family level, but some taxa were identified down to genus or species (Smith 2001, Thorp and Covich 2001, Merritt et al. 2019). Although Kerfoot et al. (2019, pers. comm., 2023) did not identify benthic copepods, we included them in our analysis.

### Zooplankton sampling and processing

Zooplankton were sampled on three dates at each beach (between June 21 and July 25, 2021) by towing a nylon plankton net (153-µm mesh; 50-cm diameter x 150-cm long opening) parallel to the shoreline along shallow (0.76–2.23 m) and deep (3.14–5.30 m) water zones. Two replicate, 5-minute tows were performed in each depth zone. Tows were 30 m behind the vessel; distance traveled (m), tow speed (cm/s), and water volume filtered (cm^3^) were recorded with a mechanical flowmeter mounted in the center of the net ring. Upon collection, the net was washed down with filtered lake water (from the outside of the net); captured zooplankton were stored in 95% ethanol and shipped to a contract laboratory for identification and enumeration (EcoAnalysts, Inc., Moscow, Idaho).

EcoAnalysts processed the zooplankton samples according to US EPA SOP LG403 Standard Operating Procedure for Zooplankton Analysis (US EPA 2016). Briefly, a sub-sample of 200–400 microcrustaceans (except nauplii) was identified to the lowest practical level and enumerated. If aquatic organisms of interest were found, 3–5 organisms of each taxon were carefully removed from the sample and isolated for further inspection and reference. Samples were identified using the most appropriate technical literature that is accepted by the taxonomic discipline and reflects the accepted nomenclature.

### Statistical analysis

All statistical analyses, including determinations of standard descriptive statistics and simple correlations (e.g., Pearson’s *r*), were performed using R (R Development Core Team 2022). We used the non-parametric Peto-Peto comparison, as implemented in the R package NADA2 (Peto and Peto 1972, Julian and Helsel 2023), to compare metal concentrations in sediments among beaches; this non-parametric method handles both non-normal data and censored data (e.g., concentration results <LOQ). The software also reports the median values for the groups, using Kaplan-Meier methods, which are appropriate for occasions where non-detects occur or distributions are not well known (Helsel 2005). For statistical comparisons of metal concentrations among sites, we focused on the 10 metals that were both quantified in most samples and were enriched at BUR-H (thought to be due to a greater relative presence of SS) relative to the other sites: Cd, Co, Cr, Cu, Fe, Mn, Ni, Pb, Se, and Zn. As previously mentioned, Tl was <LOQ at all sites; As concentrations were relatively similar among all sites (n=79; mean ± standard error [SE], 1.68 ± 0.06 mg/kg dw).

We also characterized overall sediment toxicity due to metals by comparing our total recoverable results to consensus-based probable effects concentrations (PECs; MacDonald et al. 2000). A PEC represents a concentration of a contaminant in a sediment (generally with particle size <2-mm) above which adverse effects are probable. Metals PECs are based on total concentrations (Ingersoll et al. 2000); in this way, concentrations measured following our digestion method, which generally accesses only the fraction of metal(s) that could become environmentally or biologically available, may ultimately underestimate toxicity relative to the PECs. However, total recoverable concentrations have been used previously to screen for metal hazards (e.g., Besser et al. 2015). Individual PEC values (mg/kg dw) are the following: As, 33; Cr, 111; Cu, 149; Pb, 128; Ni, 48.6; and Zn, 459. Although there is a PEC for Cd (4.98 mg/kg dw), we did not include it in our assessments because Cd concentrations were generally near or <LOQ in all sediment samples; all samples had Cd ≤0.14 mg/kg dw, so Cd contributions to overall toxicity were likely small. There is currently no PEC for Co; however, we substituted a Canadian clean-up target (80 mg/kg dw) for the purposes of our comparisons (Gray and Eppinger 2012). No PECs are available for Mn, Fe, Se, or Tl, so these metals were not included in our sediment toxicity evaluations.

We also considered PEC quotients (PEQ) to assess the potential for additive effects of multiple metal stressors. A PEQ is the ratio of the measured concentration of metal to the PEC; a PEQ quantifies the degree to which a metal concentration in a sediment sample exceeds the PEC threshold. For example, Cu has a PEC of 149 mg Cu/kg dw, so a sediment sample with 745 mg Cu/kg dw has a PEQ of 5 (745 divided by the PEC). This is somewhat analogous to the US EPA’s Hazard Quotient and Hazard Index metrics for assessments of potential risks to human health resulting from exposure to chemical mixtures (US EPA 1991). In multi-stressor situations, such as might be experienced by benthic communities at BUR due to the presence of multiple metals in the SS, overall metal hazards might usefully be estimated as ΣPEQ. In other words, a PEQ is calculated for each individual stressor to normalize the concentration to the relative toxicity of that metal; the PEQs are then summed (ΣPEQ) to generate a toxicity metric that assumes the hazards from individual metals are fully additive (Ingersoll et al. 2001, US EPA 2005, Besser et al. 2015). A ΣPEQ ≥1 is generally considered a conservative screening indicator of sediment toxicity to benthic organisms (Ingersoll et al. 2001). However, this approach does not consider potential synergistic and antagonistic effects wherein co-stressors interact to produce more-than-additive or less-than-additive toxicity (Ingersoll et al. 2001, Besser et al. 2009), nor does it consider any physical effects that SS might have on biota (such as SS smothering interstitial spaces).

To characterize the relationship between SS percentage and Cu concentration, we used a simple linear model to relate SS to Cu (as in Kerfoot et al. 2021). However, residuals from this model were non-normally distributed; therefore, we log-transformed the Cu data, which improved the distribution of residuals and other diagnostics.

We used non-metric multidimensional scaling (NMDS, with two dimensions) to visualize community composition differences (using the vegan package in R; Manly 2005). After plotting the sites in a two-dimensional space, we plotted the 95% confidence intervals and the standard deviation around the centroid for each beach. The NMDS for benthic macroinvertebrates included Gastropoda, Sphaeriidae, Hydrachnidae, Trichoptera, Nematoda, Oligochaeta, Amphipoda, Chironomidae, Copepoda, and Cladocera. The NMDS for zooplankton included Acari, Bosminidae, Calanoida, Centropagidae, Cercopagididae, Chironomidae, Chydoridae, Copepoda, Cyclopidae, Cyclopoida, Daphniidae, Diaptomidae, Ephemeroptera, Harpacticoida, Holopediidae, Ilyocryptidae, Ostracoda, Polyphemidae, Sididae, and Temoridae. Each individual was only included in the lowest taxonomic classification in which it could be identified (e.g., a Diaptomidae individual is only included in the Diaptomidae group, not the Copepoda group).

We used a generalized linear model (in base R) with beach as a categorical predictor to assess differences in benthic community composition among beach areas. This is conceptually identical to a standard analysis of variance (ANOVA) method in that it identifies whether a categorical variable is associated with changes in a continuous response variable. When using categorical predictors, one category is used as the default, and the effect of switching from that category to another category is estimated. In our models, we used BUR-H as our default category so that we could compare other beach areas to it. Standardized slopes (β) and the coefficient of determination (R^2^) values were calculated as standardized estimates of effect size (Tabachnick and Fidell 2001). Because RES samples were collected with a different sampling approach, they are not included in any of the among-beach comparisons.

We used multi-level models (R package lm4e) that included the beach as a random effect on the intercept to assess the direct relationship between SS and benthic community composition; sample sizes were too small to estimate both intercept and slope as random effects. Zooplankton data were not modeled either. Sites resampled from Kerfoot et al. (2019; RES) were treated as a separate group of sites because they represented a distinct sampling strategy. Kerfoot et al. (2019, 2021) had previously found that the model that best fit the data included both SS and SS squared (SS^2^), so we also included a model with both SS and SS^2^. The following parameters were used to represent stamp sands: SS, SS plus SS^2^, total recoverable Cu concentrations, and ΣPEQ; each of these SS parameters was included in a model that predicted either benthic invertebrate density (using a natural log distribution) or benthic invertebrate taxa count (using a Poisson distribution). Additional models included water depth, or a combination of water depth with an SS parameter. We also included a null model that included only the random effect of beach. A complete list of models (n = 12) considered is included in the results. For each model, the Akaike’s Information Criterion (corrected for small sample size; AIC_C_) value was calculated, and models were ranked according to the difference (Δ) in AIC_C_ values (lower ΔAIC_C_ values indicate a better-fit model; Burnham and Anderson 1998). Typically, ΔAIC_C_ is estimated relative to the best-fit model (i.e., the model that explains the greatest amount of variation using the fewest independent variables); models with a ΔAIC_C_ <2 are considered to have a fit equivalent to the best model, while models with ΔAIC_C_ >10 are usually considered to have much worse fit than the best model, although these are guidelines rather than objective criteria (Burnham and Anderson 1998, 2001). If two models have ΔAIC_C_ <2, but one model has more parameters than the other, the model with fewer parameters is generally considered to have stronger support (Arnold 2010). For the most strongly supported model, we estimated standardized slopes (β), and for all models we estimated marginal and conditional R^2^ values as standardized estimates of effect size (Tabachnick and Fidell 2001, Nakagawa and Schielzeth 2013, Nakagawa et al. 2017). Marginal R^2^ values indicate variation explained by the fixed effects (i.e., in this case variables such as water depth and SS), while conditional R^2^ values indicate variation explained by the marginal effects conditional on the random effects (i.e., the beach).

## Results

### Stamp sand and metals concentrations across the beach areas

Stamp sands were more prevalent at BUR-H, lower at GTB-M, and lowest at LTB-L, as evidenced by both increased percentages of SS (Fig. 2A) and greater metals concentrations in the sediment samples at BUR-H relative to the other sites (Figure 2, Table S1). We noted dark particles present in the BIG-C beach samples that were visually similar to SS. However, these particles are unlikely to be SS from the pile at Gay, Michigan, based on the relatively low concentrations of metals typically characteristic of the Gay SS (e.g., Cd, Co, Cr, Cu, Mn, Ni, Zn) relative to BUR (Figure 2); thus, we believe the black particles in BIG-C sediments were from another source (e.g., manganese sands can be confused for SS, although they are only a small fraction of the Grand Traverse Bay sands; Kerfoot et al. 2021).

**Figure 2.**
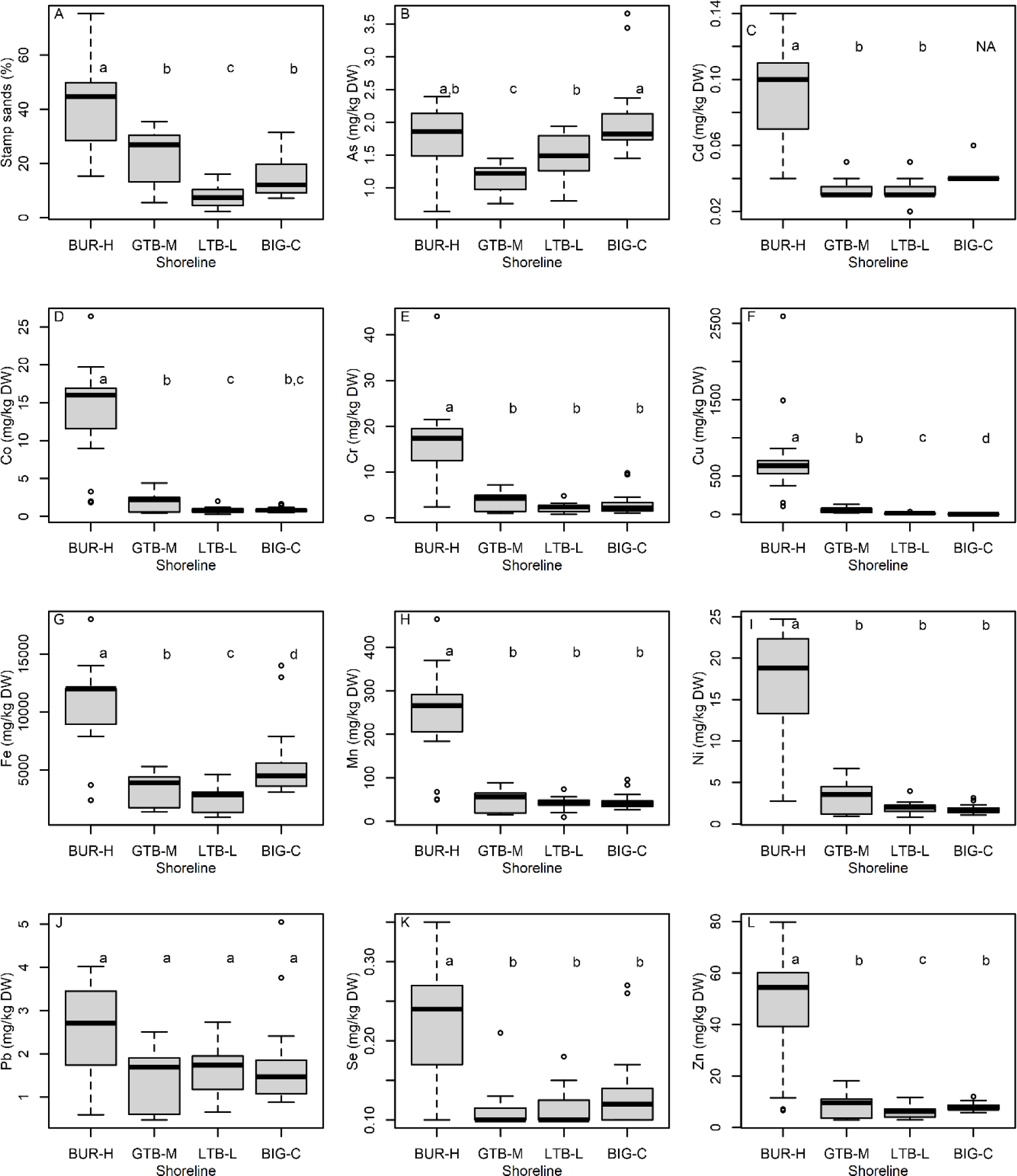
Graphical comparisons of (A) stamp sand content (percent) and total recoverable concentrations (mg/kg dry weight, DW) of (B) As, arsenic; (C) Cd, cadmium; (D) Co, cobalt; (E) Cr, chromium; (F) Cu, copper; (G) Fe, iron; (H) Mn, manganese; (I) Ni, nickel; (J) Pb, lead; (K) Se, selenium; and (L) Zn, zinc; in the sediments collected by Petite Ponar from beach sites at Buffalo Reef (BUR), Grand Traverse Bay south of Traverse River (GTB), Little Traverse Bay (LTB) and Big Bay (BIG), Michigan in 2021. Letters refer to beaches that were not clearly different using the non-parametric Peto-Peto test. In all panels, boxes encompass the first and third quartiles. The lines (whiskers) show the largest or smallest observation that falls within 1.5 times the box size. Observations that fall outside the lines are shown individually.

Sediments from the BUR-H beach were enriched in Cd, Co, Cr, Cu, Fe, Mn, Ni, Se, and Zn (Figure 2, Table S1). Individual metal PECs were only exceeded for Cu (i.e., PEQ ≥1), and then only in samples from BUR-H or RES (Figure 3), which had broad overlap with BUR-H. When PEQs were summed across 7 metals (ΣPEQ), sediment samples from 14 of 15 BUR-H sites, 13 of 16 RES sites, and 2 of 16 sites at GTB-M had a ΣPEQ ≥ 1 (Figure 3). No BIG-C or LTB-L sites had PEQs ≥1 or ΣPEQ ≥1. If we assume that estimates of SS in BIG-C sediments are zero (i.e., none of the observed black particulates were SS), the model relating SS to Cu concentrations is very strong (R^2^ = 0.84). The mean model prediction crosses the Cu PEC of 149 mg Cu/kg dw (i.e., PEQ >1) at 32% SS, although the prediction interval at this point is very wide (26–851 mg Cu/kg dw; Figure 3). Pearson’s *r* between SS and water depth is 0.42 (95% CI = 0.23–0.59); and Pearson’s *r* between ΣPEQ and water depth is 0.17 (CI = −0.06–-0.38).

**Figure 3.**
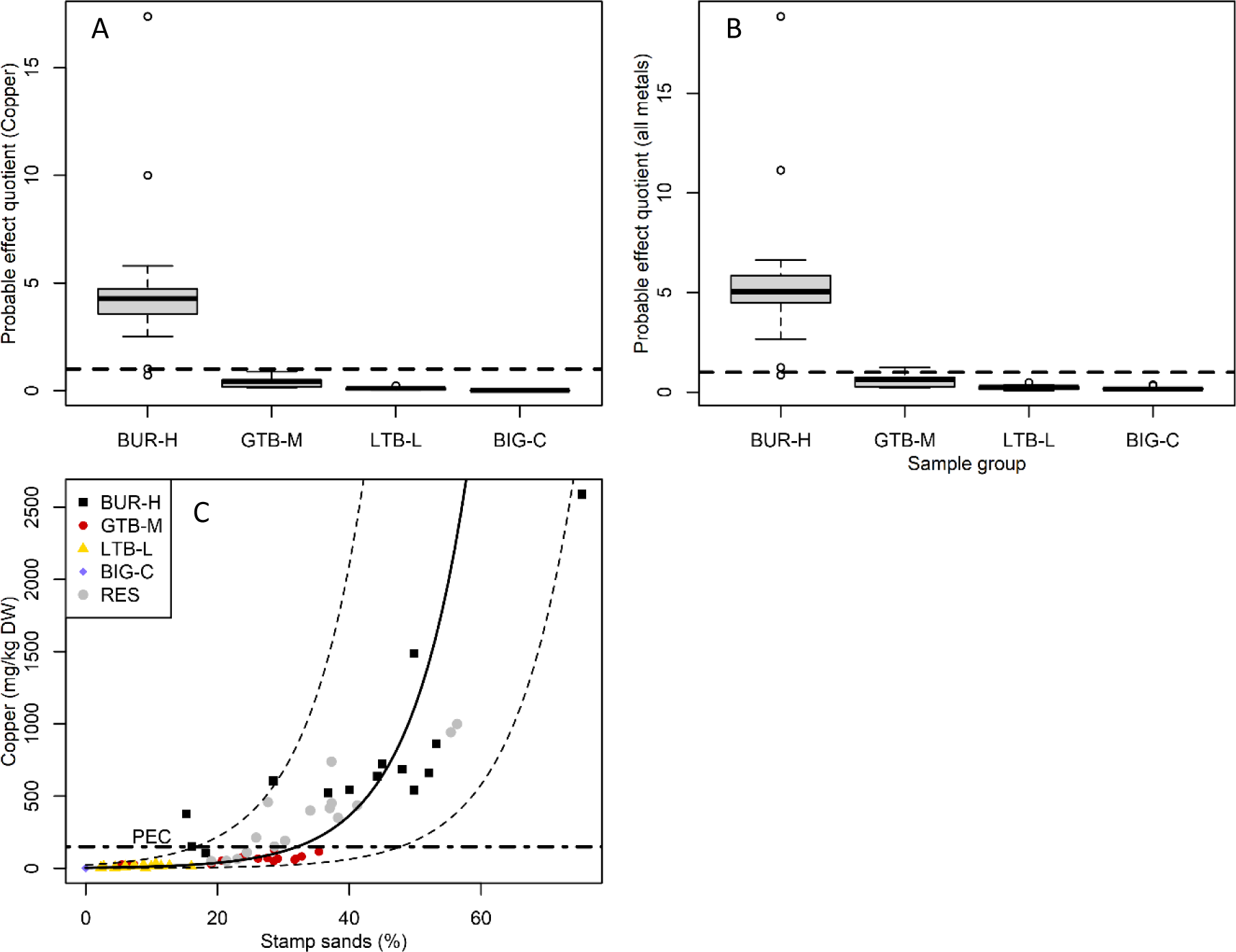
Probable effects quotients (PEQs) for total recoverable (A) copper and (B) sum of 7 metals (as ΣPEQs) in the sediments collected by Petite Ponar from study sites in the Keweenaw Peninsula, Michigan in 2021. The ΣPEQ calculations in panel B include the PEQ for copper, arsenic, chromium, lead, nickel, zinc, and cobalt; the horizontal dotted lines in panels A and B are PEQ=1, above which increased rates of adverse biological effects are expected. In panels A and B, boxes encompass the first and third quartiles. The thick black line is the median. The lines (whiskers) show the largest or smallest observation that falls within 1.5 times the box size. Observations that fall outside the lines are shown individually. (C) The bivariate relationship between stamp sands (as % of sediment) and copper concentration; the horizontal dotted line shows the probable effects concentration (PEC) for copper (149 mg/kg dw), above which biological effects are considered likely. The solid line (panel C) is the Cu concentration predicted from a model relating stamp sands to the log of the copper concentration (dashed lines indicate 95% confidence interval on the prediction). BUR-H – beach near Buffalo Reef; GTB-M – beach in Grand Traverse Bay south of the Traverse River; LTB-L – beach in Little Traverse Bay; BIG-C-beach in Big Bay (a control site); RES – samples from sites resampled from Kerfoot et al. (2019).

Particle size distributions qualitatively varied across beaches. Compared to LTB-L and BIG-C, BUR-H had a much lower proportion (%) of particles in the 125–250-µm range (medium sand) (Figure 4). BUR-H was the only beach where fine and very fine sands in the 63–125-µm range generally made up the largest proportion of the substrate composition (Figure 4). BUR-H, and RES to a lesser extent, also tended to have a greater proportion of coarse (>500 µm) particles than the other sample groups.

**Figure 4.**
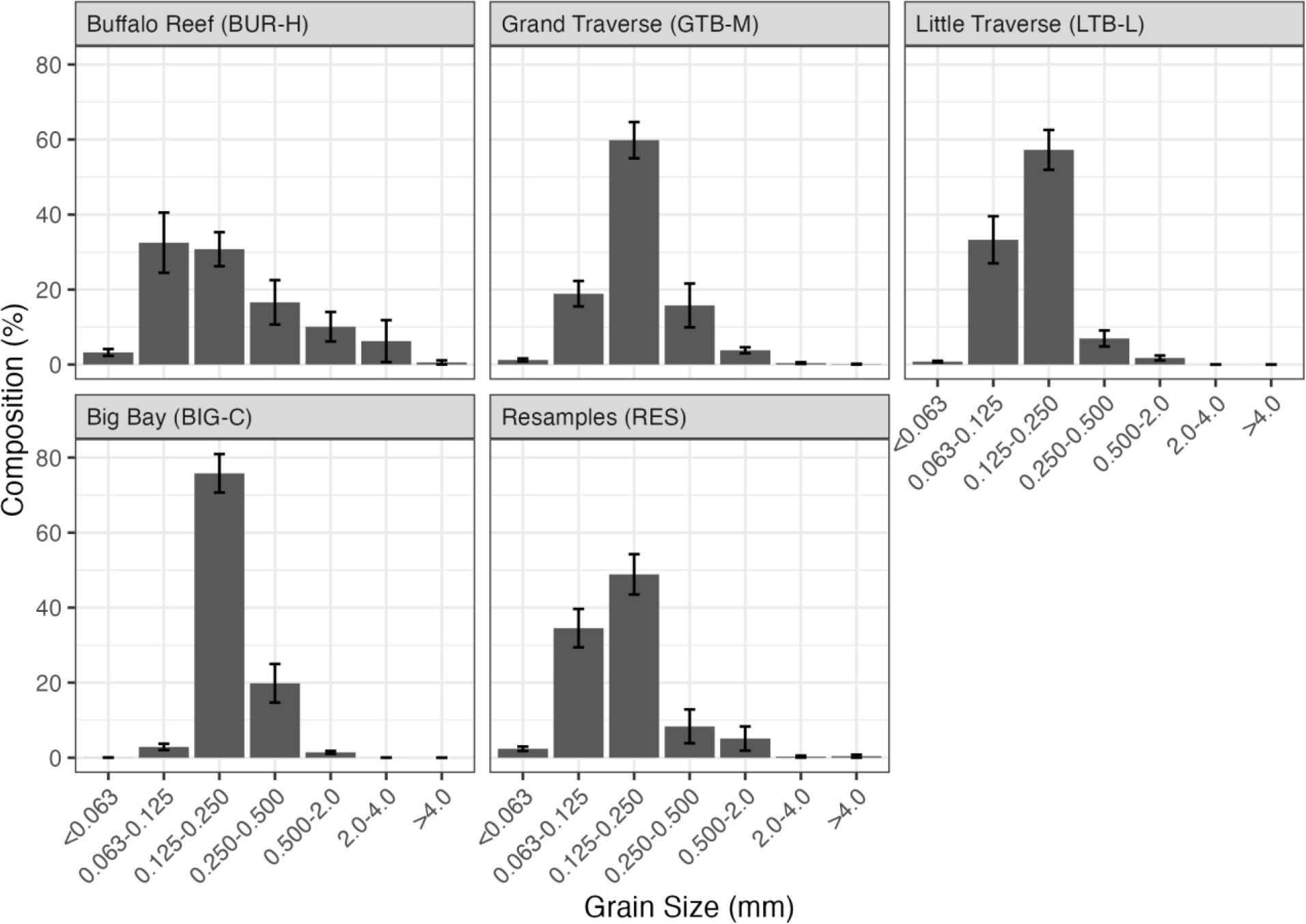
Particle size distribution in sediments collected by Petite Ponar at each beach in the Keweenaw Peninsula, Michigan in 2021. Top of bars are means and vertical lines are standard errors.

### Variation in benthic communities across beach areas

Among the benthic taxa, cladocerans, chironomids, Sphaeriidae, and oligochaetes were present at all beaches (Table 2). Nematodes, copepods, gastropods, and amphipods were absent from BUR-H, but present at one or both control sites (Table 2). Overall, the median abundance of benthic organisms was ∼3 orders of magnitude higher at LTB-L compared to other beaches (Figure 5, note log scale), driven mainly by large numbers of Harpacticoida copepods. The LTB-L beach also had the greatest number of taxa overall (Table 2, Figure 5). Looking at individual taxa, BUR-H had more *Bythotrephes longimanus* and Holopediidae than the BIG-C site, and fewer Harpacticoida, Chironomidae, Sphaeriidae, Oligochaeta, and Amphipoda than the LTB-L control site (Table 3). GTB and BUR never had clearly different numbers of these taxa (Table 3). The BUR and GTB beaches had comparable median diversity index values, whereas BIG had the lowest median diversity (Figure 5). The NMDS analyses shows that the benthic community composition at BUR-H has substantial overlap with the communities at both GTB-M and BIG-C, but very little or no overlap with LTB-L (Figure 6); in contrast, LTB-L did overlap with GTB-M and BIG-C.

**Figure 5.**
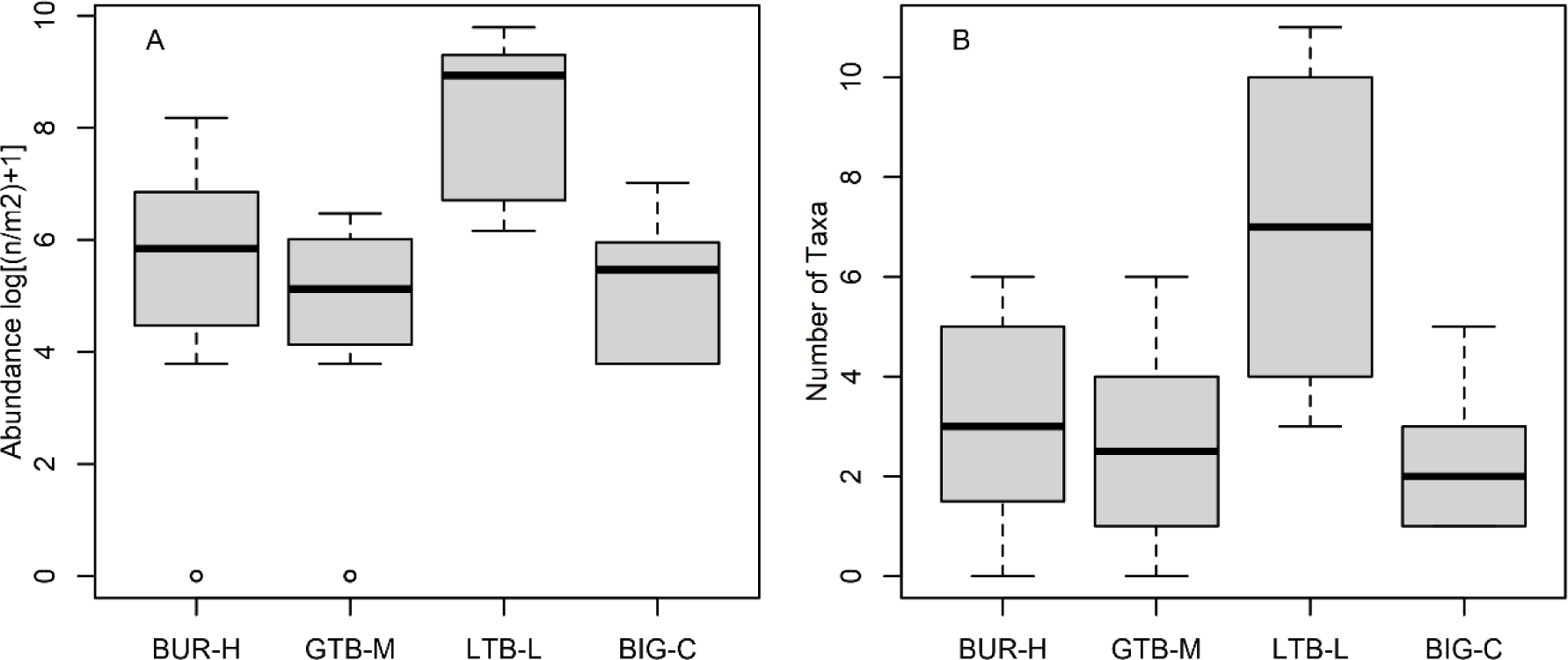
Graphical comparisons of abundance (as number, n, per square meter, m^2^) and number of taxa for benthic invertebrates collected by Petite Ponar from beach sites at Buffalo Reef (BUR-H), Grand Traverse Bay south of Traverse River (GTB-M), Little Traverse Bay (LTB-L) and Big Bay (BIG-C), Keweenaw Peninsula, Michigan, in 2021. Boxes encompass the first and third quartiles. The thick black line is the median. The lines (whiskers) show the largest or smallest observation that falls within 1.5 times the box size. Observations that fall outside the lines are shown individually.

**Figure 6.**
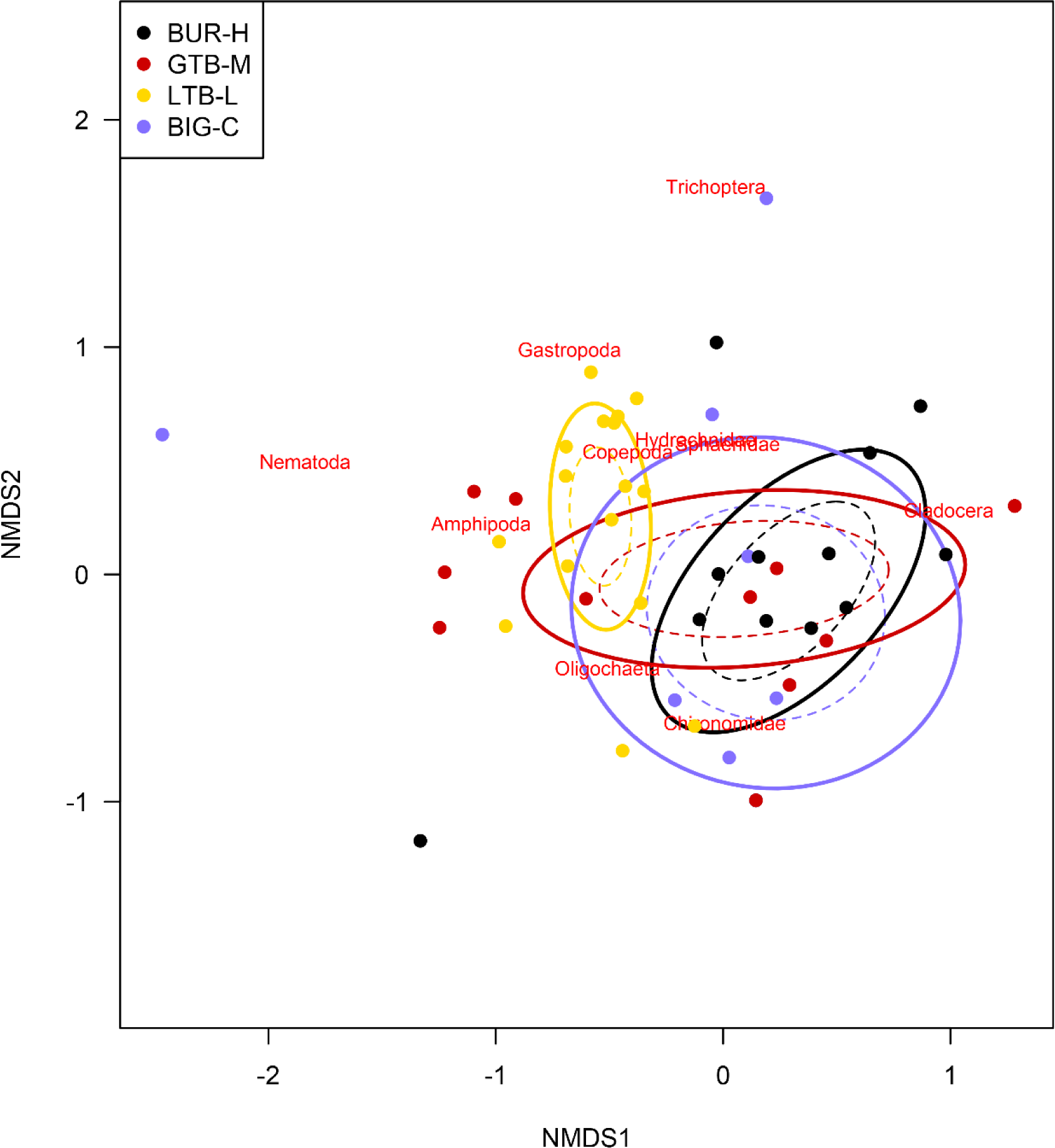
Non-metric, multi-dimensional scaling (NMDS) plot of benthic invertebrate community composition at four beaches at Buffalo Reef (BUR-H), Grand Traverse Bay south of Traverse River (GTB-M), Little Traverse Bay (LTB-L), and Big Bay (BIG-C) at the Keweenaw Peninsula, Michigan, in 2021. Solid ellipses enclose the centroid of the group within 1 standard deviation, and dashed ellipses enclose the centroid of the group with 95% confidence interval. Individual species are plotted at the point where their distribution is at a theoretical maximum. The two-dimensional stress of the NMDS ordination is 0.14.

**Table 1.** Aggregated physical and water quality characteristics of sampling areas where benthic invertebrates (all sites) and zooplankton (beach sites) were collected in 2021 to assess the impact of stamp sands (Gay, Michigan; Lake Superior).

| Sample group<sup>1</sup> | Mean Water Depth (m) | Minimum Water Depth (m)<sup>2</sup> | Maximum Water Depth (m)<sup>2</sup> | Water Temperature (°C) | Dissolved Oxygen (mg/L) | pH |
| --- | --- | --- | --- | --- | --- | --- |
| Buffalo Reef Beach (BUR-H) | 4.59 | 1.22 | 7.65 | 19.4 | 9.71 | 8.21 |
| Grand Traverse Bay Beach (GTB-M) | 4.11 | 0.73 | 8.20 | 19.0 | 9.18 | 8.21 |
| Little Traverse Bay Beach (LTB-L) | 2.55 | 0.61 | 4.66 | 18.9 | 9.28 | 8.15 |
| Big Bay Beach (BIG-C) | 3.65 | 0.61 | 7.65 | 19.8 | 8.74 | 8.23 |
| Resampled sites (RES, in Grand Traverse Bay) | 7.64 | 5.21 | 9.48 | 18.8 | 9.86 | 8.28 |

<sup>1</sup> Resampled sites include 16 sites that were previously sampled in Kerfoot et al. (2019). All other beaches were sampled along four transects, each consisting of four individual locations, across a gradient from shallow to deep water.
<sup>2</sup> Water depths at the shallowest and deepest points along the transects are reflected as the minimum and maximum depths, respectively.

**Table 2.** Benthic taxa observed at benthic sampling locations in the Keweenaw Peninsula, Michigan, 2021.

| Order/Phylum | Buffalo Reef<br>(BUR-H) | Grand<br>Traverse Bay<br>(GTB-M) | Little<br>Traverse Bay<br>(LTB-L) | Big Bay<br>(BIG-C) | Sites resampled<br>from Kerfoot et<br>al. (2019; RES) |
| --- | --- | --- | --- | --- | --- |
| Cladocera (other than those<br>identified below) | X | X | X | X | X |
| • *Bythotrephes longimanus* | X | X | X | X | X |
| • Holopediidae spp. | X | X | X | X | X |
| Harpacticoida |  |  | X | X |  |
| Cyclopoida |  | X |  |  | X |
| Calanoida |  |  |  |  | X |
| Diptera (Chironomidae) | X | X | X | X | X |
| Trichoptera | X |  | X | X | X |
| Gastropoda |  |  | X |  |  |
| Sphaeriidae | X | X | X | X | X |
| Oligochaeta | X | X | X | X | X |
| Amphipoda (*Diporeia*) |  | X | X |  | X |
| Hydrachnidia | X |  | X |  |  |
| Nematoda |  | X | X | X | X |
| Total Orders/Phyla | 6 | 7 | 10 | 7 | 9 |

**Table 3.**
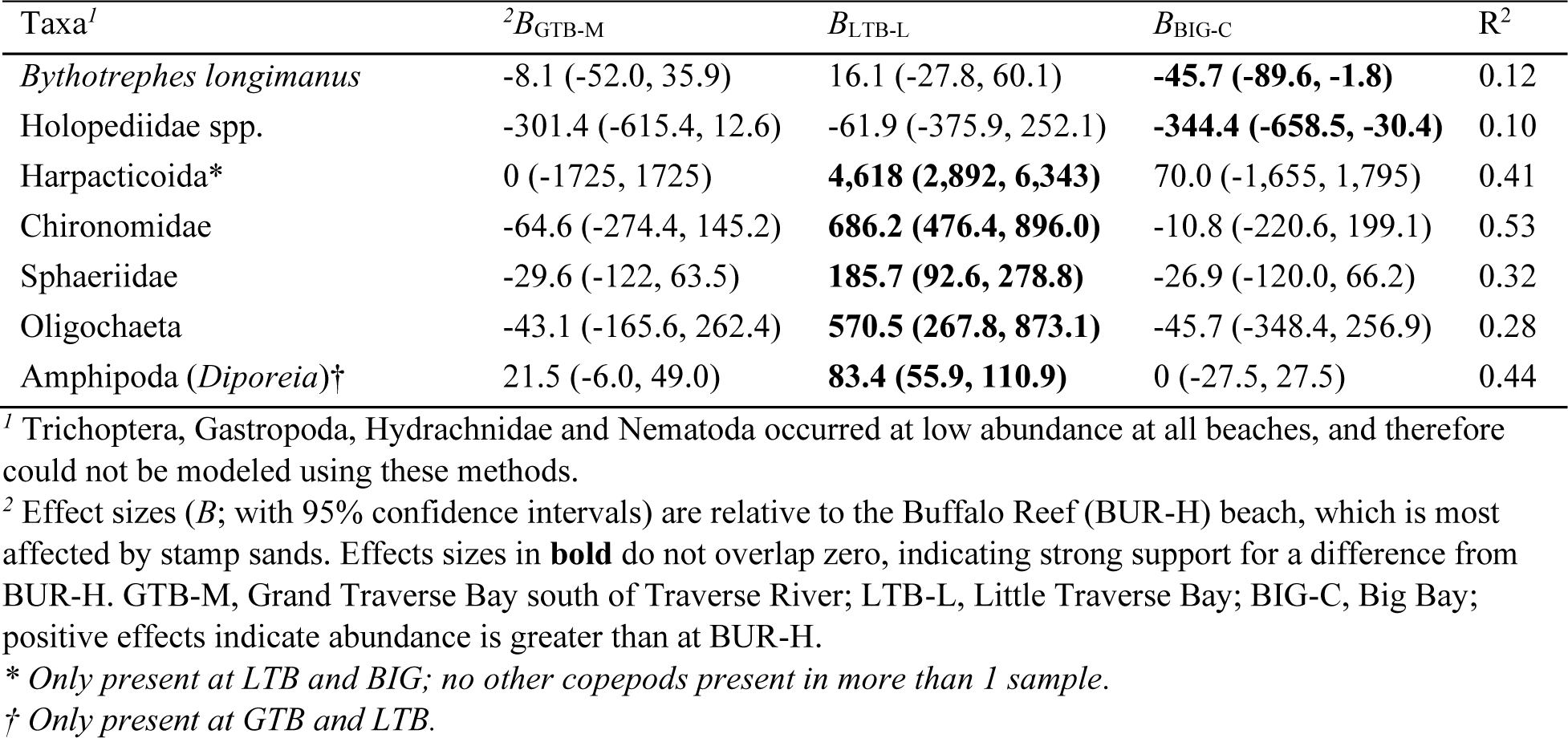
Differences in zooplankton density (log count per m^3^) among beaches, estimated using a generalized linear model at sites in the Keweenaw Peninsula, Michigan, 2021.

### Variation in zooplankton communities across beach areas

Twenty taxa of invertebrates were identified in the zooplankton samples. Many juvenile copepods were only identified as copepods or to order, but four families of copepods were identified (Centropagidae, Diaptomidae, Temoridae, Cyclopidae; Table 4). Cladocerans included eight families; most other taxa were only identified to sub-class or order (Table 4). A few typically benthic taxa were observed in the plankton nets (e.g., Ephemeroptera once, Chironomidae 10 times; Table 4) and were included in our analyses of the zooplankton. We did not exclude these taxa because they do occur in the pelagic zone at times (e.g., while moving or during disturbance) and because excluding them has no significant effect on the results. The total number of zooplankton taxa observed was greatest at LTB-L (Table 4), where 18 total taxa were identified), whereas other areas had lower diversity (12 taxa at BUR-H, 11 at GTB-M, and 13 at BIG-C). However, individual samples showed considerable variation within sites (Figure 7). Three families of Cladocerans (Ilyocryptidae, Polyphemidae and Sididae), insect taxa (Chironomidae and Ephemeroptera), Acari, and Harpacticoida were absent at BUR-H (Table 4). Zooplankton NMDS plots show considerable overlap between the zooplankton community compositions at BUR-H and GTB-M, but LTB-L and BIG-C were very distinct from the two other beaches (Figure 8).

**Figure 7.**
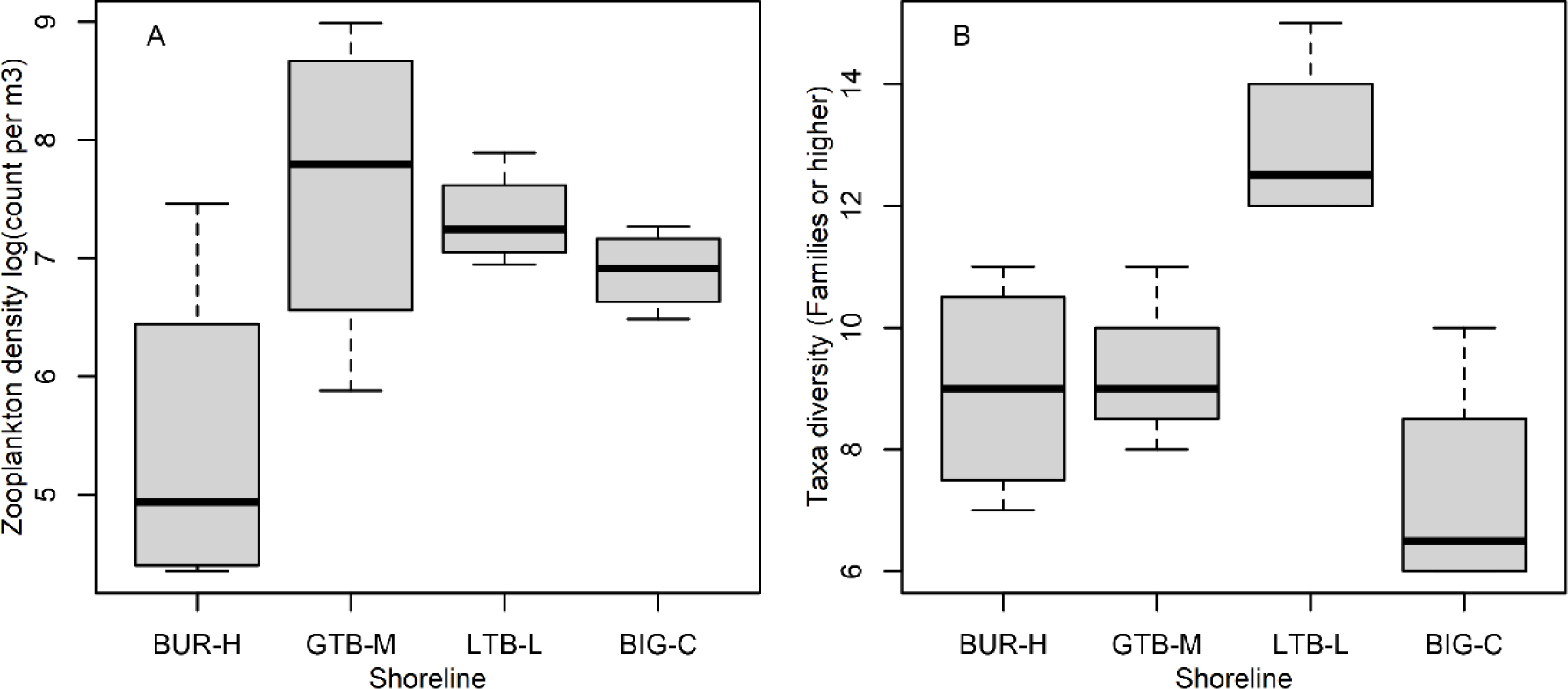
Graphical comparisons of zooplankton density (count per cubic meter, m^3^) (A) and diversity (B) by beach at Buffalo Reef (BUR-H), Grand Traverse Bay south of Traverse River (GTB-M), Little Traverse Bay (LTB-L) and Big Bay (BIG-C) on the Keweenaw Peninsula, 2021. Boxes encompass the first and third quartiles. The thick black line is the median. The lines (whiskers) show the largest or smallest observation that falls within 1.5 times the box size.

**Figure 8.**
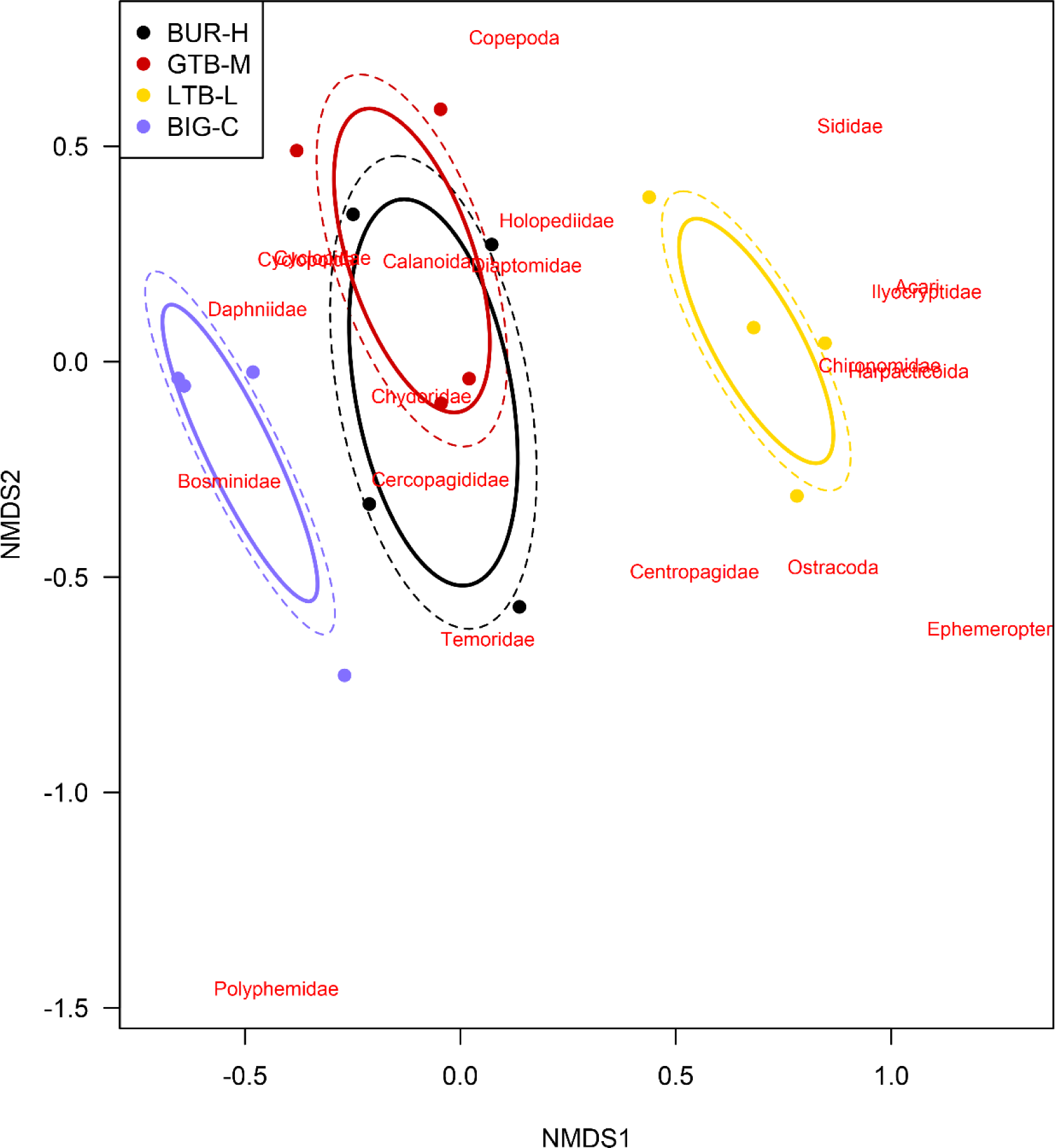
Non-metric, multi-dimensional scaling (NMDS) plot of zooplankton community composition at four beaches in Lake Superior: Buffalo Reef (BUR), Grand Traverse Bay south of Traverse River (GTB), Little Traverse Bay (LTB), and Big Bay (BIG) on the Keweenaw Peninsula, Michigan in 2021. Solid ellipses enclose the centroid of the group within 1 standard deviation, and dashed ellipses enclose the centroid of the group with 95% confidence interval. Individual species are plotted at the point where their distribution is at a theoretical maximum. The order Cyclopoida and the family Cyclopidae (within the same order) appear in the exact same space on this NMDS plot (which obscured the labeling); therefore, only Cyclopoida is labeled. The two-dimensional stress of the NMDS ordination is 0.17.

**Table 4.**
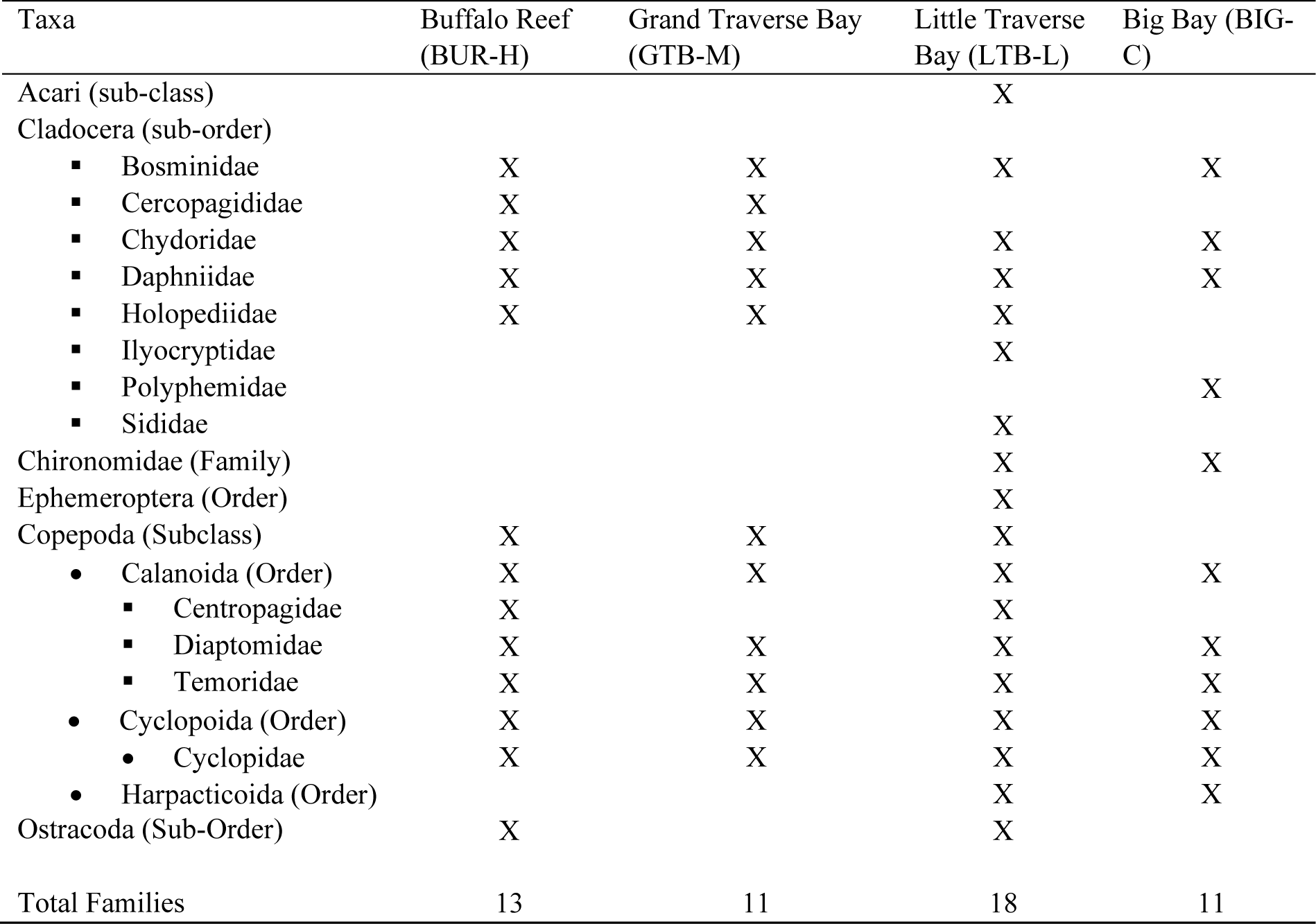
Zooplankton taxa observed at the benthic sampling locations in the Keweenaw Peninsula, Michigan, 2021.

Models indicated that total zooplankton (density) was 1–2 orders of magnitude greater at GTB-M, LTB-L and BIG-C relative to BUR-H (Table 5), which is also apparent from the data (Figure 7). Nine taxa (Cladocera and Copepoda) were common to at least three beaches (Table 4), none of which were clearly denser at BUR-H than at other sites (Table 5). As an overall group, Cladocera were more common at all three of the less impacted sites (GTB-M, LTB-L, and BIG-C) than BUR-H, but individual taxa varied (Table 5). For example, Daphniidae were more common at GTB-M than BUR-H, but LTB-L and BIG-C were not clearly distinct from BUR-H. For Copepoda as a group, GTB-M and LTB-L had higher densities than BUR-H, but BIG-C was not clearly distinct (Table 5). Again, when looking at lower taxonomic levels these patterns differed. For example, all of the sites had similar numbers of Temoridae (Table 5).

**Table 5.** Differences in zooplankton density (log count per m^3^) among beaches, estimated using a generalized linear model in the Keweenaw Peninsula, Michigan, 2021.

| Taxa | <sup>1</sup>*B*<sub>GTB-M</sub> | *B*<sub>LTB-L</sub> | *B*<sub>BIG-C</sub> | R<sup>2</sup> |
| --- | --- | --- | --- | --- |
| Total Zooplankton | **2.2 (0.8, 3.6)** | **1.9 (0.5, 3.3)** | **1.5 (0.1, 2.9)** | 0.48 |
| Cladocera | **1.1 (0.1, 2.2)** | **2.1 (1.1, 3.1)** | **2.9 (1.9, 3.9)** | 0.74 |
| ▪ Bosminidae | 0.2 (-1.2, 1.6) | 0.3 (-1.1, 1.8) | **3.3 (1.9, 4.8)** | 0.70 |
| ▪ Chydoridae* | 0.3 (-0.9, 1.4) | **3.5 (2.3, 4.6)** | **2.4 (1.2, 3.5)** | 0.81 |
| ▪ Daphniidae | **1.6 (0.05, 3.1)** | 0.4 (-1.1, 2.0) | 1.1 (-0.4, 2.7) | 0.29 |
| ▪ Holopediidae† | **2.6 (1.8, 3.4)** | **3.1 (2.2, 3.9)** | -0.7 (-1.6, 0.1) | 0.91 |
| Copepoda | **2.3 (0.8, 3.7)** | **1.5 (0.03, 3.0)** | 1.2 (-0.2, 2.7) | 0.44 |
| • Calanoida | **1.4 (0.3, 2.5)** | **1.5 (0.4, 2.5)** | -0.7 (-1.8, 0.3) | 0.66 |
| ▪ Diaptomidae* | 1.4 (-0.1, 2.9) | **2.3 (0.8, 3.8)** | 0.5 (-1.0, 2.0) | 0.48 |
| ▪ Temoridae | 0.5 (-0.4, 1.4) | 0.4 (-0.5, 1.3) | 0.3 (-0.6, 1.1) | 0.11 |
| • Cyclopoida | **3.1 (1.2, 5.0)** | 1.8 (-0.1, 3.7) | **2.3 (0.4, 4.2)** | 0.47 |
| ▪ Cyclopidae | **3.0 (1.4, 4.5)** | 1.1 (-0.5, 2.7) | 1.3 (-0.3, 2.8) | 0.53 |

<sup>1</sup> Effect sizes (*B*; with 95% confidence intervals) are relative to the Buffalo Reef (BUR-H) beach, which is most affected by stamp sands. Effects sizes in **bold** do not overlap zero, indicating strong support for a difference from BUR-H. GTB-M, Grand Traverse Bay south of Traverse River; LTb-L, Little Traverse Bay; BIG-C, Big Bay; positive effect indicate the abundance is greater than BUR-H.
\* *Not present at BUR.*
† *Not present at BIG.*

### Direct relationships between invertebrates and stamp sand abundance

We used multi-level models to examine direct relationships between invertebrates and SS (i.e., SS, Cu concentrations, and ΣPEQ). For model estimation, SS at the BIG sites were set to zero. We also included data from the 16 sites resampled from Kerfoot et al. (2019; RES sites). Each of the supported models for both density and richness included water depth (i.e., all models with ΔAIC_C_ <10), which had a strong positive effect on invertebrate density and richness (Table 6, Figure 9A, D). The strongest model for benthic density included both water depth and Cu concentration; this model had an R^2^ of 0.73, with a marginal R^2^ of 0.28 (Table 6). Similarly, the strongest model for taxonomic richness had an R^2^ of 0.66 and a marginal R^2^ of 0.32 (Table 6). We note that although models with water depth and Cu were ranked highest by ΔAIC_C_, models with ΣPEQ instead of Cu had very similar support (Table 6). Both Cu and ΣPEQ had negative effects on both benthic density and taxonomic richness in these models (Figure 9B, C, E, F; Table 6). Because ΣPEQ is a composite variable representing the overall sediment toxicity experienced by the benthic community (including Cu), parsimony indicates ΣPEQ should be conceptually weighted more heavily than Cu alone (Burnham and Anderson 1998). Models with only SS had much less support, although the second order polynomial regression (the model with a SS^2^ term) did fit the data better than a null model for both benthic density and taxonomic richness (Table 6).

**Figure 9.**
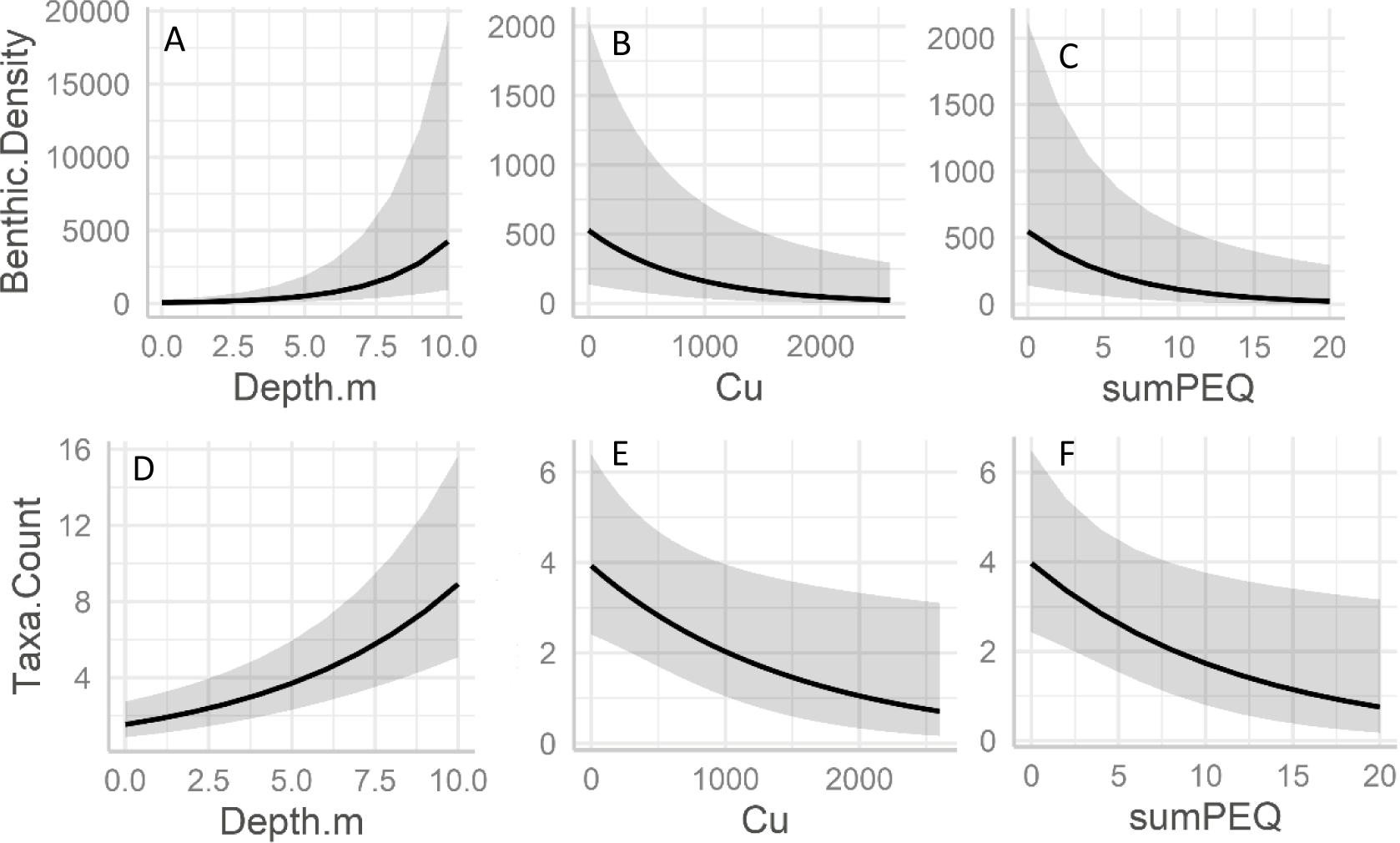
Model predictions of conditional effects of water depth (m), copper (Cu) concentration in the sediment (mg/kg dry weight), and the sum of probable effect quotients (ΣPEQ, denoted as sumPEQ in the figure) from seven metals in the sediment on benthic density (individuals per m^2^; A,B,C) and diversity (taxa count; D,E,F) in samples collected from Grand Traverse Bay, Little Traverse Bay and Big Bay, Keweenaw Peninsula, 2021. Model predictions are conditional on the beach area being sampled (in this case the Buffalo Reef beach). Two models are depicted for each response variable, one that includes Cu concentration (B, E) and one that includes ΣPEQ (C, F). The effect of water depth is identical in both models(A,D). These correspond to the two most strongly supported models in Table 6. The solid line is the predicted mean, and the grey shading encompasses the mean ± standard error of the prediction.

**Table 6.** Models relating stamp sand (SS) content, copper concentration in the sediment (Cu), the sum of the probable effects quotients (ΣPEQ) for 7 metals in the sediment, and water depth (Depth) to the density and taxonomic richness of the benthic invertebrate community in the Keweenaw Peninsula, Michigan, 2021.

| Response | Model<sup>1</sup> [with $\beta^2$ (confidence interval)] | ΔAIC<sub>C</sub> | R<sup>2</sup><sub>Marginal</sub> | R<sup>2</sup><sub>Conditional</sub> | Standardized intercepts ($\alpha$) |
| --- | --- | --- | --- | --- | --- |
| Benthic density | **Cu × -0.24 (-0.43, -0.06) +**<br>**Depth × 0.57 (0.39, 0.75)** | **0** | **0.28** | **0.73** | **$\alpha_{BUR} = 0.10$; $\alpha_{GTB} = -0.76$;**<br>**$\alpha_{LTB} = 1.34$; $\alpha_{BIG} = -0.37$; $\alpha_{RES} = -0.34$** |
| [log(n m<sup>-2</sup>+1)] | ΣPEQ × -0.25 (-0.43, -0.06) +<br>Depth × 0.59 (0.40, 0.75) | 0.1 | 0.28 | 0.73 | $\alpha_{BUR} = 0.10$; $\alpha_{GTB} = -0.77$;<br>$\alpha_{LTB} = 1.34$; $\alpha_{BIG} = -0.37$; $\alpha_{RES} = -0.33$ |
|  | Depth | 4.3 | 0.27 | 0.73 |  |
|  | SS + SS<sup>2</sup> + Depth | 5.7 | 0.30 | 0.75 |  |
|  | SS + Depth | 6.0 | 0.27 | 0.71 |  |
|  | SS + SS<sup>2</sup> | 26.4 | 0.22 | 0.67 |  |
|  | Cu | 30.1 | 0.10 | 0.48 |  |
| | ΣPEQ | 31.0 | 0.09 | 0.48 | |
|  | Null | 35.2 | - | 0.39 |  |
|  | SS | 37.3 | <0.01 | 0.40 |  |
| Taxonomic Richness | **Cu × -0.27 (-0.52, -0.04) +**<br>**Depth × 0.47 (0.30, 0.64)** | **0** | **0.32** | **0.66** | **$\alpha_{BUR} = 1.36$; $\alpha_{GTB} = 0.92$;**<br>**$\alpha_{LTB} = 1.98$; $\alpha_{BIG} = 0.78$; $\alpha_{RES} = 1.07$** |
| Species # | ΣPEQ × -0.25 (-0.49, -0.04) +<br>Depth × 0.47 (0.30, 0.65) | 0.3 | 0.31 | 0.66 | $\alpha_{BUR} = 1.36$; $\alpha_{GTB} = 0.92$;<br>$\alpha_{LTB} = 1.98$; $\alpha_{BIG} = 0.78$; $\alpha_{RES} = 1.07$ |
|  | SS + SS<sup>2</sup> + Depth | 3.1 | 0.34 | 0.67 |  |
|  | Depth | 3.5 | 0.29 | 0.67 |  |
|  | SS + Depth | 3.5 | 0.28 | 0.65 |  |
|  | SS + SS<sup>2</sup> | 23.2 | 0.24 | 0.54 |  |
|  | Cu | 28.1 | 0.13 | 0.49 |  |
| | ΣPEQ | 29.3 | 0.11 | 0.47 | |
|  | Null | 33.4 | - | - |  |
|  | SS | 35.6 | <0.01 | 0.38 |  |

<sup>1</sup> All models were multi-level models that included a beach grouping variable. Taxonomic richness models used a Poisson distribution because species counts are discrete. Data that did not come from the regular beach sampling (i.e., RES sites resampled from Kerfoot et al. 2019) were in a separate grouping variable.
<sup>2</sup> All slope ( $\beta$ ) estimates are standardized.
The most strongly supported model is highlighted in **bold**.

## Discussion and Conclusions

We observed distinct invertebrate communities in areas directly impacted by SS, as did Kerfoot et al. (2019, 2021). SS covered BUR-H and this site was depauperate in benthic taxa that are important prey items for the development of native fishes, such as *Diporeia* and benthic copepods; moreover, zooplankton density was orders of magnitude lower at BUR-H than at the other beaches. Although SS have multiple mechanisms to impair invertebrate taxa in the benthos (e.g., toxicity from metals, abrasion, physical blockage of habitat), the concentration of Cu (or ΣPEQ, the combination of Cu and other metals sourced from the SS) was more strongly associated with declines in benthic abundance and diversity than was SS in the most strongly supported models. This indicates that exposure to Cu in the SS (and the resulting uptake and toxicity) was the primary cause of the observed effects on macroinvertebrates; the stark effects on zooplankton (discussed further below) are additional evidence that Cu toxicity is the major contributor to effects of SS on the invertebrate community at BUR-H. Other metals may also contribute some to the overall toxicity, but the evidence in our study did not demonstrate that, as no other metal ever exceeded its PEC and including other metals in a predictive model did not improve the fit.

Kerfoot et al. (2021) used a second order polynomial to model the association between SS and invertebrate community structure and diversity. Here we used a log-normal model structure (by using a log transformation) to model associations between measurements of SS and invertebrate structure and diversity. Kerfoot et al. (2021) noted that this non-linearity indicates that the toxicity of SS varies across the range of SS percentages. Kerfoot et al. (2021) also observed that SS that have traveled farther appear to have lower Cu concentration. As lower concentrations of SS correspond to sites farther from the original SS source, this would explain why relationships between SS and Cu or SS and invertebrate community directly are non-linear. However, our model was non-linear even when using Cu data directly. This implies that the effect of Cu and other metals themselves are non-linear and synergistic effects are likely occurring in the field.

Unlike benthic taxa, zooplankton are unlikely to be affected by physical abrasion or filling of interstitial spaces in the sediment. This means SS effects are likely driven by metal leaching from sediments either directly (via toxicity) or indirectly (via food web effects). Relative to other sites, BUR-H had fewer zooplankton cladocerans and copepods (up to 2 orders of magnitude fewer). This varied among the individual taxa within those larger groups, but as a food resource for larval fish, the zooplankton community as a whole was clearly affected at BUR-H. Cladocerans and copepods were the most abundant zooplankton taxa, and copepods were the most abundant benthic invertebrate. We know that not only are copepods and other aquatic invertebrates and vertebrates sensitive to dissolved Cu in the water column (e.g., Flemming and Trevors 1989, Heuschele et al. 2022), but that waterborne Cu can result in major changes in the planktonic community structure (Banerjee et al. 2021). We do not have metals data for water to demonstrate potential leaching of Cu into the water column from the SS at our study sites. However, we believe it is reasonable to assume that this is the source of reduced abundance at BUR-H. Numerous invertebrate taxa are sensitive to Cu exposure (e.g., Besser et al. 2018), either directly or via trophic transfer mechanisms (e.g., Campana et al. 2012). Water-only metals exposures have previously been shown to poorly predict biological effects from sediment toxicity, including that of sediments collected in the Keweenaw Waterway and elsewhere (West et al. 1993, de Castro-Català et al. 2016), but the data here indicate these biological effects can be severe for the zooplankton community’s overall abundance.

Our data, although limited to a single year, indicates that literature-based definitions of sediment toxicity (i.e., exceedances of PECs, ΣPEQ ≥1) may not be protective of community structure and diversity in these beach areas. Most BUR-H sites had Cu concentrations above the PEC, while all of the GTB-M and LTB-L sites had concentrations below the Cu PEC. However, the overall picture from the invertebrate data here indicates BUR-H and GTB-M are more similar to each other than to LTB-L. We hypothesize that GTB-M is already experiencing effects from SS-derived Cu, even though this site has concentrations below the PEC threshold, and therefore Cu concentrations would need to be kept lower than the PEC to prevent negative impacts effects. In addition to the PEC, MacDonald et al. (2000) also reported consensus-based threshold effect concentrations (TECs), below which harmful effects are unlikely to be observed. The TEC value for Cu (31.6 mg/kg dw) was exceeded in 100% of the BUR-H and RES samples, as well as in 11 of the GTB samples (69%) and 1 LTB (6%) sample. The Cu TEC threshold may be a better indicator of effects in our study data than the PEC, with GTB-M and BUR-H on one side of the TEC and LTB-L on the other, consistent with the invertebrate community data. Although our data are consistent with GTB having already suffered impacts from Cu derived from SS, we lack a before-after dataset that would make a compelling case for this. The possibility exists that LTB-L has always been very different than GTB-M and BUR-H, even prior to SS arrival.

Predicting and modeling Cu from SS concentrations is challenging. Concentrations of Cu in the Gay, Michigan, SS have been measured in several studies, but the exact leaching and weathering processes that occur and potentially contribute to biological effects once the sands enter and drift within Lake Superior are largely unknown. Soil sampling (i.e., onshore sampling of SS deposit areas) by the Michigan Department of Environmental Quality; MDEQ 2006) indicated a mean Cu concentration of 2,683 mg/kg (range 1,500–13,000) in soils collected from areas closest to the conveyor that was used to transport sands into Lake Superior at the time of mill operation. If we were to extrapolate using the mean Cu concentration from those undisturbed soils, then this would indicate that Cu PECs would be exceeded when SS are more than about 5% (range 1–10%). This is less than the lowest observed value where Cu PEC was exceeded in our dataset (15.3%), indicting the SS we observed had lower Cu content than the original pile. Another deposit area, which represented an onshore area in which SS that had eroded from the main pile were deposited, had a mean Cu concentration of 1,443 mg/kg (range 710–5,300; MDEQ 2006). These SS had experienced more movement and weathering, and either these processes had leached Cu from the SS or particles with less Cu are more mobile (Kerfoot et al. 2021). These weathered SS would need to make up 10% (3–21%) of the sands at a site to reach the Cu PEC, which is closer to the low end of our observations. To our knowledge, this is the first time that SS have been observed to make up >10% of the sand at any LTB-L sites; indeed, we observed SS >10% a total of 4 times. Some of these LTB-L sands could be manganese sands previously discussed in Kerfoot et al. (2021), but in Kerfoot et al., the dark manganese sands made up, on average, just 1.8% of grain counts. Given the relatively low concentrations of SS required to reach the Cu PEC and given the Cu PEC may not be protective of the invertebrate communities, this may indicate that the LTB-L site may be in imminent danger of experiencing impacts to the invertebrate community.

Our results indicate that continued direct measurements of Cu and other metals in SS collected from Lake Superior would be important because extrapolation of Cu from SS collected from previous onshore soil studies could misrepresent *in situ* conditions for effects determinations and remedial purposes (Kerfoot et al. 2021). Additionally, SS heterogeneity within each beach likely results in localized exposure conditions to fish eggs, larvae, and the benthos. In other words, the benthos, including larval fish, could be exposed to locally elevated metals concentrations due to differing SS within their home ranges, based on relative immobility compared to adult fish. Localized effects are supported by our models, which indicated that Cu concentrations in the sediments and water depth were more critical for benthic density and taxonomic richness than SS.

Ultimately, our observations of reduced benthic abundance and taxa at BUR relative to beaches farther afield from Gay, Michigan, indicate that SS inundation has resulted in biological and community effects. Our models indicate that >15% SS is associated with sediment toxicity due to Cu; many of our GTB-M sites have reached this threshold. Additional studies of SS toxicity, both in terms of SS thresholds (particularly given the possibility that SS have begun reaching LTB-L) and associated biological effects to the benthos and larval fish are warranted. Land managers could consider differences in sensitivities among different life stages and taxa (e.g., de Castro-Català et al. 2016) and use multi-pronged biological and chemical investigations, including quantification of sediment toxicity modifiers (e.g., organic carbon, particle size; Roman et al. 2007, Campana et al. 2012) to (1) establish site-specific sediment quality assessment frameworks for the Keweenaw Peninsula, and (2) evaluate remedial options to restore the critical nursery habitat at Buffalo Reef. Larval fish raised on Buffalo Reef and growing on nearby beaches are likely to be affected by the declines in early life-stage food resources (e.g., zooplankton) that we observed here.

## Acknowledgements

Any use of trade, firm, or product names is for descriptive purposes only and does not imply endorsement by the U.S. Government. Thank you to two U.S. Geological Survey researchers who provided peer review on an earlier version of this manuscript.

## Data availability statement

Data used in this study can be accessed at https://doi.org/10.5066/P13DWTRR (Larson et al., 2024).

---

**Table S1.** Comparison of metal concentrations among four beaches potentially impacted by a stamp sands (SS) source near Gay, MI. The most highly impacted beach is near Buffalo Reef (BUR-H). Another beach in Grand Traverse Bay south of the Traverse River is moderately impacted by SS (GTB-M) and very low SS are believed to occur at a beach in nearby Little Traverse Bay (LTB-L). We also sampled a control beach, Big Bay (BIG-C). We used the Peto-Peto test, which is a non-parametric test appropriate when some data are below the quantification limit and the data are non-normally distributed. Medians (standard deviation) are in units of mg/kg dw and were estimated using Kaplan-Meier methods to account for occasions when values were below the limit of quantification (Helsel 2005). The Peto-Peto test indicated concentration differences between beaches having different letters.

|  | BUR-H | GTB-M | LTB-L | BIG-C |
| --- | --- | --- | --- | --- |
| Arsenic | 1.74 (0.58) a,b | 1.16 (0.21) c | 1.48 (0.38) b | 2.07 (0.62) a |
| Cadmium | 0.09 (0.03) a | 0.03 (0.01) b | 0.03 (0.01) b | Below LOQ† |
| Cobalt | 13.9 (7.0) a | 1.98 (1.22) b | 0.85 (0.42) c | 0.88 (0.32) b, c |
| Chromium | 16.4 (10.1) a | 3.73 (1.99) b | 2.29 (1.02) b | 3.16 (2.69) b |
| Copper | 745 (601) a | 60.1 (32.8) b | 14.0 (8.4) c | 1.70 (0.30) d |
| Iron (x 10<sup>3</sup>) | 10 (4) a | 3.3 (1.4) b | 2.5 (1.1) c | 5.7 (3.3) d |
| Lead | 2.60 (1.20) a | 1.40 (0.70) a | 1.60 (0.60) a | 1.80 (1.10) a |
| Manganese | 243 (116) a | 48.2 (25.4) b | 41.6 (25.4) b | 45.6 (19.5) b |
| Nickel | 16.5 (7.8) a | 3.30 (1.80) b | 2.00 (1.80) b | 1.80 (0.58) b |
| Selenium | 0.22 (0.07) a | 0.11 (0.03) b | 0.11 (0.02) b | 0.14 (0.05) b |
| Zinc | 47.1 (22.8) a | 9.00 (4.70) b | 6.00 (2.30) c | 7.90 (1.70) b |

† A single BIG site had a positive detection.

